# Effect of chromatin flexibility on diffusive loop extrusion via molecular slip-links

**DOI:** 10.1101/275214

**Authors:** C. A. Brackley, J. Johnson, D. Michieletto, D. Marenduzzo

## Abstract

We use Brownian dynamics simulations to study the formation of chromatin loops through diffusive sliding of molecular slip links, mimicking the behaviour of cohesin-like molecules. We recently proposed that diffusive sliding is sufficient to explain the extrusion of chromatin loops of hundreds of kilo-base-pairs (kbp), which may then be stabilised by interactions between cohesin and CTCF proteins. Here we show that the elasticity of the chromatin fibre strongly affects this dynamical process, and find that diffusive loop extrusion is more efficient on stiffer chromatin regions. Efficiency is also enhanced if cohesin loading sites are close to regions where CTCF is bound. In light of the heterogeneous physical properties of eukaryotic chromatin, we suggest that our results should be relevant to the looping and organisation of interphase chromosomes *in vivo*.

## Introduction

Chromosome conformation captures (3C) techniques, and their high-throughput variant, so-called “Hi-C”, have provided an enormous amount of data on the 3-dimensional organisation of chromosomes of different organisms, and of different cell types in the same organism [1–3]. The holy grail of these studies is to establish the relationship between such 3-D structure and function, or gene expression.

Hi-C experiments have shown that the genomes of a number of organisms are organised into domains, called “topologically-associating domains”, or TADs [3]. In mammals (but not in bacteria, yeast or the fly), an important class of TADs are those enclosed within a loop bringing together binding sites of the CTCF proteins [4–6]. The contacts are such that the base of the loop provides the TAD boundaries. Here, CTCF is likely bound to cohesin [7–11], a ring-like protein which is thought to be able to loop chromatin, either by dimerising and forming a “hand-cuff” (Fig. 1A(i)) [10], or as a single ring enclosing two fibres [7, 10] (Fig. 1A(ii,iii)).

**Fig. 1:**
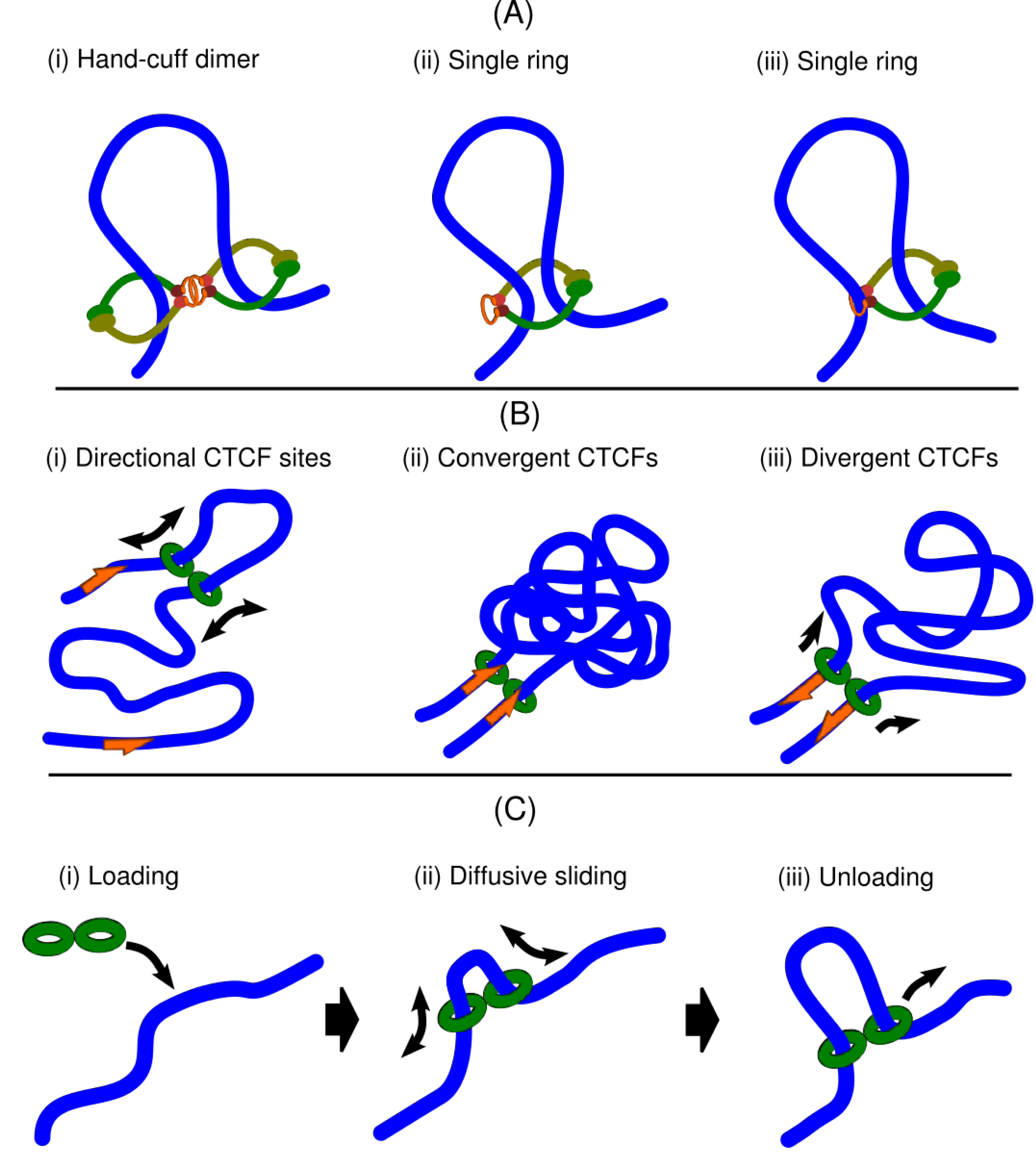
Schematic diagram illustrating the background and setup of our model. (A) Possible models for a cohesin-like molecule bound to a chromatin fibre (blue). In (i) two cohesin rings form a dimer, with the topology of a hand-cuff; in (ii) and (iii) a single cohesin ring is embracing two chromatin fibres – the difference is that in (ii) the fibres enter the same pore in the ring, whereas in (iii) they occupy different pores. While for concreteness in what follows we focus on case (i), the results do not depend on the microscopic model assumed – all that is needed is that cohesin has the topology of a slip link which can either move actively or diffuse along the chromatin fibre. (B) Diagram illustrating the bias, found in Hi-C, favouring the formation of loops between convergent CTCF binding sites. (C) Illustration of our model: a cohesin dimer is loaded on the fibre (i), after this a chromatin loop grows and shrinks as the two rings in the dimer diffuse (ii), and finally the dimer detaches (is unloaded) from the fibre (iii).

As the binding sites for CTCF are non-palindromic, they have an orientation along the chromatin (Fig. 1B(i)). Strikingly, Hi-C revealed that in the vast majority of cases (*>* 90% the two CTCF binding sites at the base of the loops have a convergent orientation (Fig. 1B(ii)) [2]; only a few have parallel orientations, and virtually none a divergent orientation (Fig. 1B(iii)). A possible explanation for this seemingly puzzling bias for convergent loop formation was suggested in Refs. [12, 13], which generalised a biophysical model previously introduced in Ref. [14]. These authors proposed a loop extrusion model according to which cohesin (or another bivalent loop-extruding factor) is able to bind chromatin and actively move along the fibre, in such a way that the genomic distance between the segments brought together by the hand-cuff grows linearly in time. [The idea that a ring-like protein, such as condensin or cohesin, may work as a motor driving loop extrusion dates back to Ref. [15].] If loop extrusion is halted when cohesin meets a CTCF whose binding site is oriented towards it (but continues through CTCFs oriented the other way), then the convergent bias is naturally explained.

A concern with this active loop extrusion model is, however, that a loop-extruding factor with a motor activity has yet to be found. Whilst condensin has recently been shown to be able to move unidirectionally on DNA [16] (with the direction presumably selected by spontaneous symmetry breaking), experiments with cohesin have thus far only reported diffusive sliding [17–19], and never active unidirectional motion. But is a motor really necessary to explain the convergent loop bias? We recently showed that it is not [20, 21], and proposed an alternative model of diffusive loop extrusion (Fig. 1C), where a cohesin dimer binds to the chromatin fibre (Fig. 1C(i)) and diffuses (Fig. 1C(ii)) until it either unbinds (Fig. 1C(iii)) or sticks to a bound CTCF protein (Fig. 1B(ii)). Just as in the active loop extrusion model we additionally assumed that the CTCF-cohesin interaction depends on their relative orientation (cohesin just diffuses away again if the CTCF is pointing away from it). Another possible way to dispense with an explicit motor activity of cohesin has recently been proposed in Ref. [22], where the authors suggest that supercoiling generated by transcription is sufficient to power the extrusion process.

Diffusive loop extrusion can lead to the formation of a 100-kbp convergent CTCF loop within *∼* 20 min (the measured cohesin residence time on chromatin [17–19]) if the diffusion of cohesin on chromatin is 10 kbp^2^/s or more, which appears to be possible given current *in vitro* measurements. For instance, acetylated cohesin was reported to diffuse at 0.1 *μ*m^2^s on reconstituted chromatin [19], and assuming a compaction of 20 bp/nm on the fibre, which is relevant for an open 10-nm fibre *in vivo* [23, 24], we infer a diffusion coefficient of 40 kbp^2^/s. [Active extrusion, on the other hand, would require looping factors or cohesin to move at a speed of about 5 kbp/min.]

Here, we further characterise the diffusive sliding of cohesin on chromatin fibres of different stiffness by means of Brownian dynamics simulations. While the study in Ref. [20] mainly focussed on the case of a flexible fibre, it is of interest to see how the results differ when the chromatin is stiffer. This is because the persistence length of chromatin *in vivo* cannot be easily measured directly, and values estimated experimentally range between 40 and 200 nm [25]. It is typically assumed that the the lower and upper end of this range correspond to euchromatin and heterochromatin, respectively. The former being transcriptionally active and more swollen, while the latter being transcriptionally silent and more compact [26].

We find that the flexibility of the underlying chromatin fibre plays a major role in determining the efficiency of diffusive sliding of cohesin in creating large loops (which can then be stabilised by binding to convergent CTCF pairs). When the chromatin fibre is stiff, we find that diffusing molecular slip-links mimicking cohesins travel much farther with respect to the case where chromatin is flexible. The efficiency of diffusive loop extrusion therefore increases substantially with chromatin stiffness. We speculate that this may facilitate the formation of large loops on inert chromatin regions (here defined as void of active or inactive histone marks). These regions are normally assumed to be associated with linker histone H1 [27], which microscopy suggests can stiffen the fibre locally.

## Model and methods

Here, we briefly describe our simulation method. More details are given in Ref. [20] (see in particular the corresponding Supporting Information).

We perform Brownian dynamics simulations of a chromatin fibre, modelled as a beadand-spring polymer (with *N* = 2000 beads, each of size *σ*), where beads are strung together by finite-extension-nonlinear-elastic (FENE) bonds (see, e.g., [20]). A key role in this work is played by the persistence length, which determines the fibre stiffness, and which is introduced through a Kratky-Porod potential, defined in terms of the positions of a triplet of neighbouring beads along the polymer as follows:

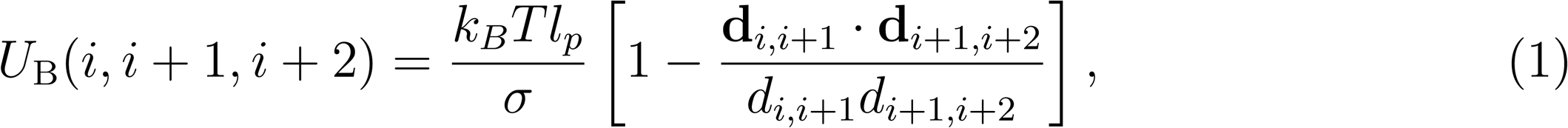

where we denote the position of the centre of the *i*-th chromatin bead by **r**_*i*_, the separation vector between beads *i* and *j* by **d**_*i,j*_ = **r**_*i*_ − **r**_*j*_, and its modulus by *d*_*i,j*_ = |**r**_*i*_ − **r**_*j*_|. Other contributions to the polymeric force field are as in Ref. [20]. CTCF binding sites are modelled as stretches of 6 beads on the polymer which are placed every 100 beads; we assume that each stretch models a pair of binding sites, and that slip-links (cohesin dimers) bind strongly to the first bead in a stretch facing them, so as to give a directionality to the binding sites and form convergent loops.

Molecular slip-links, which simulate cohesin dimers, are modelled as two rigid rings. Each of these is composed of 12 beads, arranged in a square (with side 4*σ*), with an additional phantom sphere at the centre which interacts only with beads on the chromatin fibre modelling CTCF binding sites. The two rings are held together by a pair of FENE bonds, and they are kept in an open “handcuff” arrangement via two sufficiently strong bending interactions (the potential has the same functional form as in Eq. (1)). The rototranslational motion of the centre of mass of the ring is described by suitable Langevin equations (see [20] for more details).

The slip-link beads interact with each other, and with chromatin beads with a Weeks-Chandler-Anderson potential (see Ref. [20]). The CTCF-cohesin interaction is modelled via a Lennard-Jones potential between the first bead in a CTCF stretch and the phantom bead in the middle of the slip-link rings (see above): the maximal value of this potential is 17.7 *k*_*B*_*T* and its range is 1.8 *σ*.

Previously [20] we considered two different cases: in the first, each time a slip-link is attached to the chromatin a random location on the fibre is chosen; in the second, the slip-links can only attach at special beads, or “loading sites”. Here we focus on the second case, which is relevant since *in vivo* the cohesin-loading factor (NIPBL in humans, or Scc2 in yeast) binds at preferred genomic locations, and there is some evidence that cohesin is loaded near the promoters of active genes [11]. In the simulations cohesin attachment is achieved by first positioning the slip-link in a folded handcuff arrangement such that each ring encircles an adjacent polymer bead; the bending interactions between the two rings then act to open the the handcuff, and bend the polymer inyo a loop. After this the slip-link is free to diffuse in 3-D and along the polymer, growing and shrinking the loop. Attachment is attempted at a rate *k*_on_, and occurs provided a configuration without steric clash can be found. Detachment from the fibre occurs at a rate *k*_off_: when in the detached state the position of the beads in the handcuffs is not explicitly considered in the simulation.

The chromatin fibre and slip-links are enclosed in a cubic simulation box (size 200*σ*, so that the polymer is in the dilute regime).

The mapping from simulation to physical units can be made as follows. Energies are mapped in a straightforward way as they are measured in units of *k*_*B*_*T*. To map length scales from simulation to physical units, we set the diameter,*σ*, of each bead to, for instance,*∼* 15 nm ≃ 1 kbp (assuming a chromatin fibre with compaction intermediate between a 10 nm and a 30 nm fibre; of course, all of our results would remain qualitatively valid with a different mapping). The *l*_*p*_ values we consider are between 2*σ* and 10*σ* (see snapshots in Fig. 2), hence they may be mapped to *∼* 30 − 150 nm. These values are reasonable for active euchromatin and inactive heterochromatin respectively [25]. For time scales, we need to estimate the typical diffusive timescale (over which a bead diffuses a distance comparable to its own size), or Brownian time, which equals *τ*_*B*_ ≡*σ*^*2*^*/D*. One way to do this is to require that the mean square displacement of a polymer bead matches that of a chromatin segment measured *in vivo* in Ref. [28]. This is similar to the scheme used in Refs. [20, 29], and it should be noted that, in this way, we match the effective *in vivo* viscosity, hence effectively take into account any macromolecular crowding within the nucleoplasm. We obtain *τ*_*B*_ =*σ*^*2*^*/D* ≃ 0.01 s, whereas one simulation time step is 0.01*τ*_*B*_. The on and off rate for slip-link/chromatin binding were set to 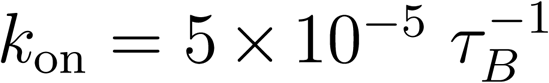 and 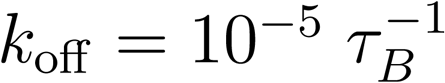 respectively, whereas simulations were run for 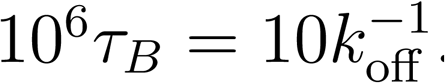. When both rings in a slip-link are bound to CTCF, we assumed that the off-rate decreased to 0, to model CTCF-induced stabilisation of cohesin-mediated off chromatin loops (similar results would be found for a decrease of the off-rate to 10-fold or more, as 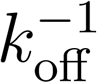 would then equal or exceed our simulation time).

**Fig. 2:**
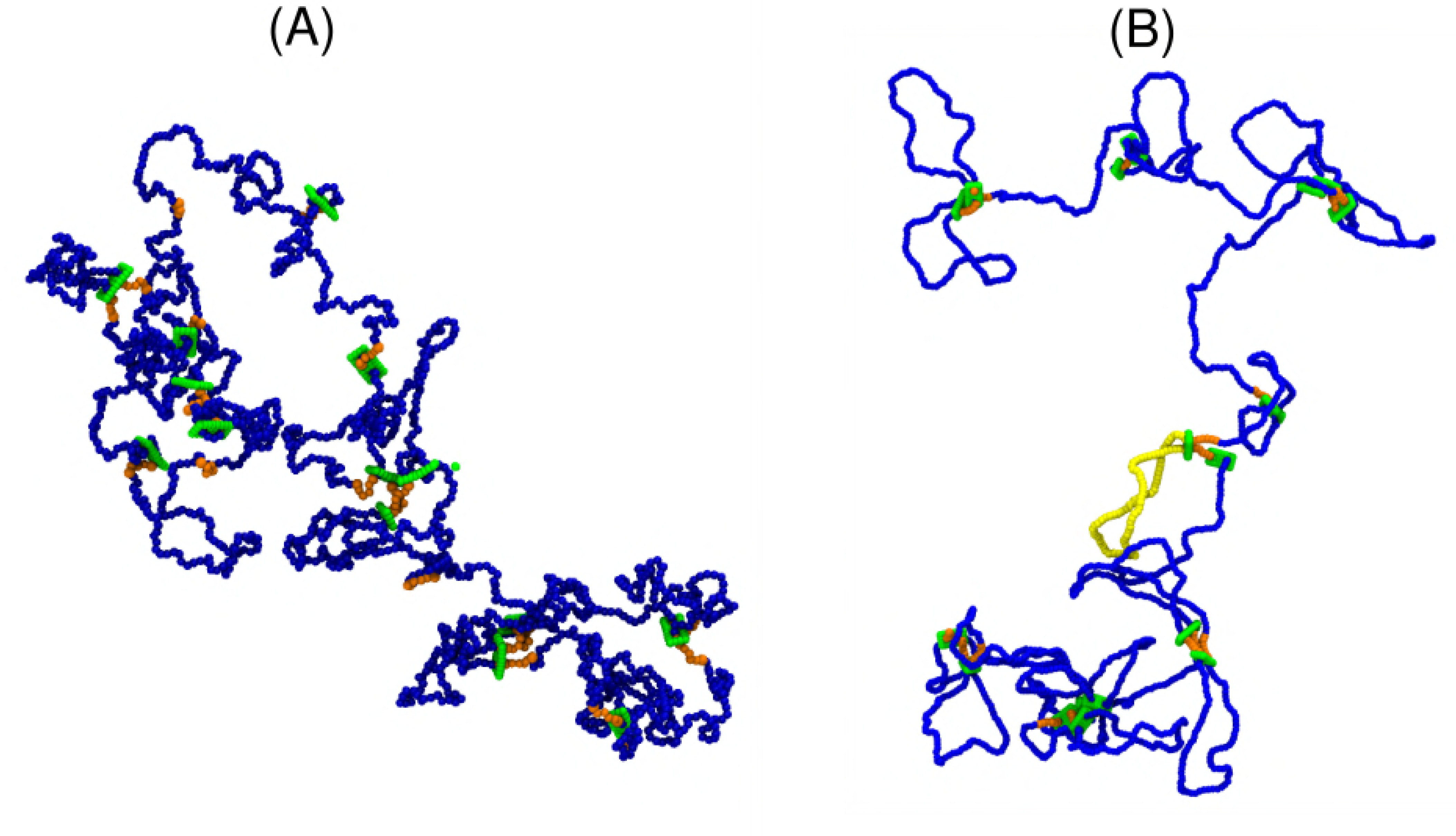
Snapshots from 3-D simulations of (A) a flexible (persistence length 2*σ*) and (B) a semi-flexible (persistence length 10*σ*) chromatin fibre. Blue and orange beads correspond to standard chromatin beads and CTCF binding sites respectively, whereas beads making up slip-links are depicted in green. In (B), one complete chromatin loop (between two neighbouring CTCF binding sites) is highlighted in yellow.

## Results

We first analyse the diffusive sliding of cohesin-like slip-links on a chromatin fibre of uniform stiffness and contour length 2 Mbp (corresponding to 2000 beads). The model chromatin is split up into sections of 100 kbp (100 beads). At the ends of each section we place a convergent pair of CTCF binding sites (see Model and Methods). Molecular slip-links attach at loader sites located in the middle of each section, and detach with uniform probability from any position on the fibre. We add 20 slip-links – for simplicity we associate each with a different section (so there are never multiple slip-link per section in this case).

Figure 3 shows the position versus time of each cohesin monomer in selected slip-links, for a flexible (persistence length 2*σ*, Fig. 3A) and a stiff (persistence length 10*σ*, Fig. 3B) chromatin fibre. The trajectories show that diffusive sliding can create large loops. Such trajectories are unlike those of standard random walks, but are instead characterised by many short excursions and a few larger ones, some of which lead to successful CTCF loop formation. As we shall see, this is because the entropic cost of looping acts to limit loop size, and can be qualitatively modelled as a confining potential.

**Fig. 3:**
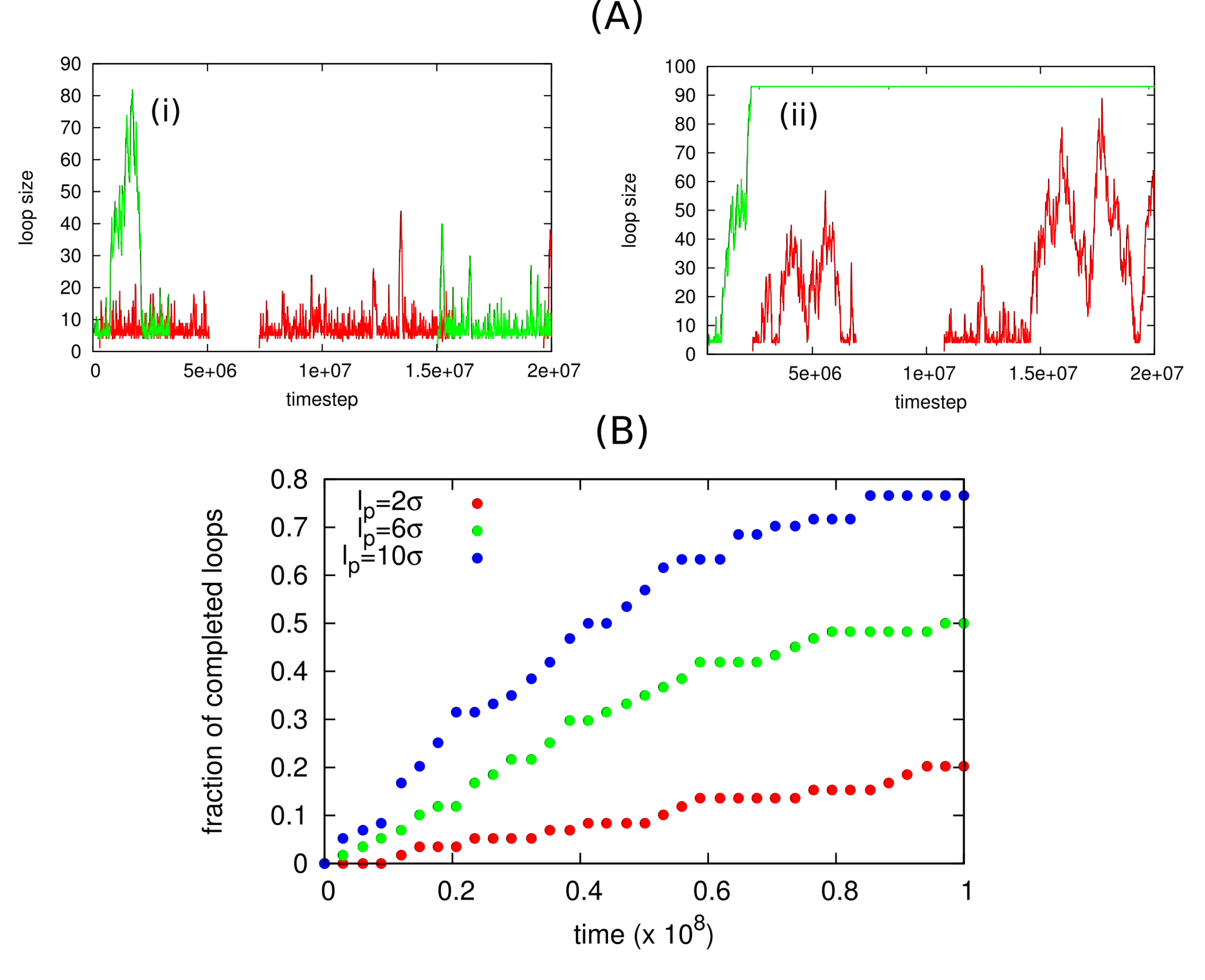
Results from 3-D simulations of a chromatin fibre split into sections by CTCF sites, with a single slip-link per section. (A) Example of loop size as a function of time for two selected slip-links (green and red curves) on a (i) flexible (*l*_*p*_ = 2*σ*) and (ii) stiff (*l*_*p*_ = 10*σ*) chromatin fibre. (B) Percentage of completed convergent CTCF loops as a function of time, for *l*_*p*_ = 2 *σ, l_p_* = 6*σ*, and *l*_*p*_ = 10*σ*.

Inspection of the trajectories also suggests that diffusive loop extrusion is more efficient on the stiffer substrate. Quantitatively, we found that at any given time the fraction of convergent CTCF loops created by diffusive sliding is a lot larger on the stiff fibre (Fig. 3C). The extent of the effect is perhaps surprising, given the factor of 5 difference between the stiff and the flexible fibres in our simulations.

To understand why chromatin flexibility affects slip-link diffusivity, we analyse a simple 1-D model of a random walker (slip-link) injected at a loading site, and diffusing in an effective potential modelling the entropic and enthalpic cost associated with looping of a semi-flexible polymer. The position of the random walker represents the size of a slip-link loop (i.e., the separation of the two side of the link). A suitable effective potential (defined up to an irrelevant additive constant), *V*, is the following [20, 30]

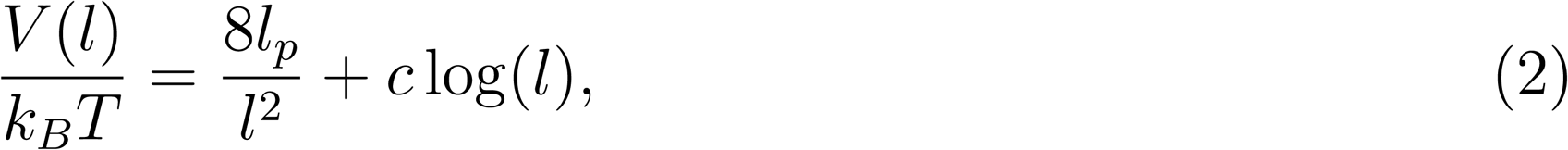

where *l* is the loop size, or position of the random walker, *l*_*p*_ is the persistence length, and *c* a universal exponent describing the entropic cost of looping (for phantom polymers without excluded volume, *c* = 3/2 in 3-D). This functional form captures the competition between the bending energy cost, which decreases monotonically with loop size *l*, and the entropic cost, which increases with *l*. For an ideal flexible polymer, the minimum of the potential will therefore be at 0 (in practice, though, this case is of limited interest as self-avoidance alone is sufficient to create a non-zero effective bending rigidity).

In the 1-D model, the random walker moves within a domain of size *L*, representing a chromatin section flanked by convergent CTCF sites as in our 3-D simulations. The random walker has a uniform probability of unbinding from the chromatin fibre, unless it has reached the CTCF pair (its position reaches *L*), in which case, for simplicity, we assume it forms a permanent loop (i.e., these configurations are absorbing states). Even without the absorbing state, this would be a non-equilibrium model because the binding and unbinding rates violate detailed balance. This is appropriate for cohesin/chromatin interactions, as both binding and unbinding are ATP-dependent, hence are active processes.

By simulating this simple 1-D problem, we can find the probability that a CTCF loop has formed as a function of time after attachment – the associated curve is plotted in Figure 4 for different values of the bending rigidity. As the chromatin stiffness contribution favours loop enlargement when the loop is small, we find that the 1-D model qualitatively reproduces the bias in favour of larger loops for stiffer fibres which is observed in the 3-D simulations (Fig. 4).

**Fig. 4:**
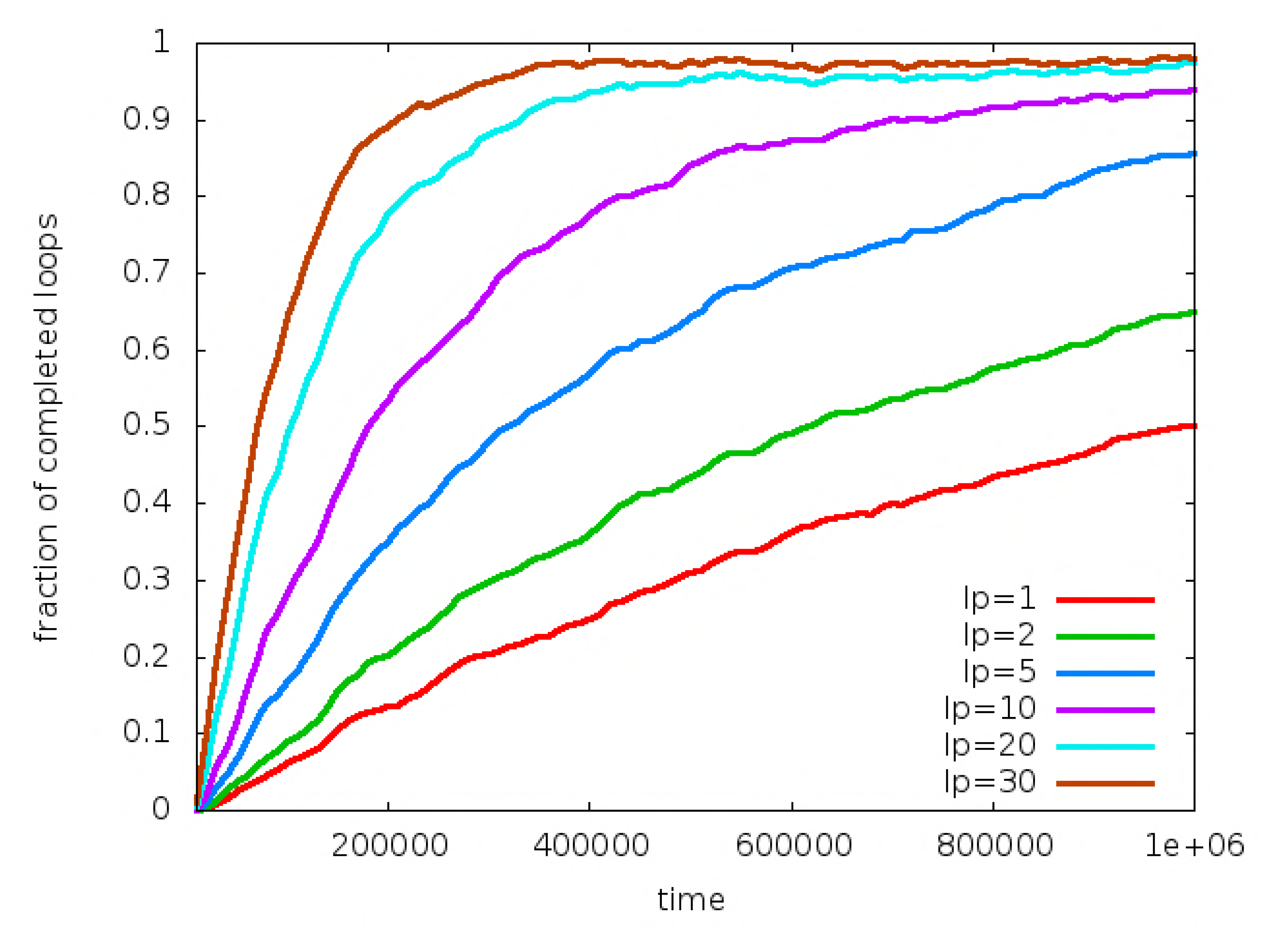
Results from simulations of the 1-D model. Percentage of 100 kbp loops formed for different values of *l*_*P*_ (given in kbp in the legend), as predicted via simulations of the 1-D model defined by the potential in Eq. (2). The random walk starts at *l* = 1 and the detachment rate *k*_off_ is set to 0 for simplicity. Assuming a baseline cohesin diffusivity (for motion with no potential) of 200 kbp^2^/s leads to a mapping of 1 time simulation units to 0.01 s (the typical residence time of cohesin on chromatin *in vivo* is *∼* 20 min [10], or *∼* 1.2 × 10^5^ simulation time units).

An important parameter in our model is the number of cohesins (slip-links) which should be present in each convergent CTCF domain. The copy number of cohesin is not known with high precision, partly because it is not straightforward to single out chromatin-bound cohesin. As is the case for other intracellular proteins, copy number may also differ in different cell types.

The simulations in Figures 3 and 4 correspond to a situation where there is one slip-link per 100 kbp (this corresponds to about 60, 000 molecules in a diploid nucleus). In Figure 5 we return to our 3-D simulations and consider the effect of increasing slip-link number, ranging from 1 to 5 per domain. We simulate a block copolymer with alternating stiff and flexible domains (see snapshots in Fig. 5A), so that the effect of different stiffness can be examined in a single simulation. Also, we placed the slip-link loader close to one of the CTCF sites; this is likely to be more realistic biologically, as cohesin loading is thought to occur close to promoters, which broadly correlate with open chromatin, DNase-hypersensitive sites and strong CTCF binding sites [3, 6].

**Fig. 5:**
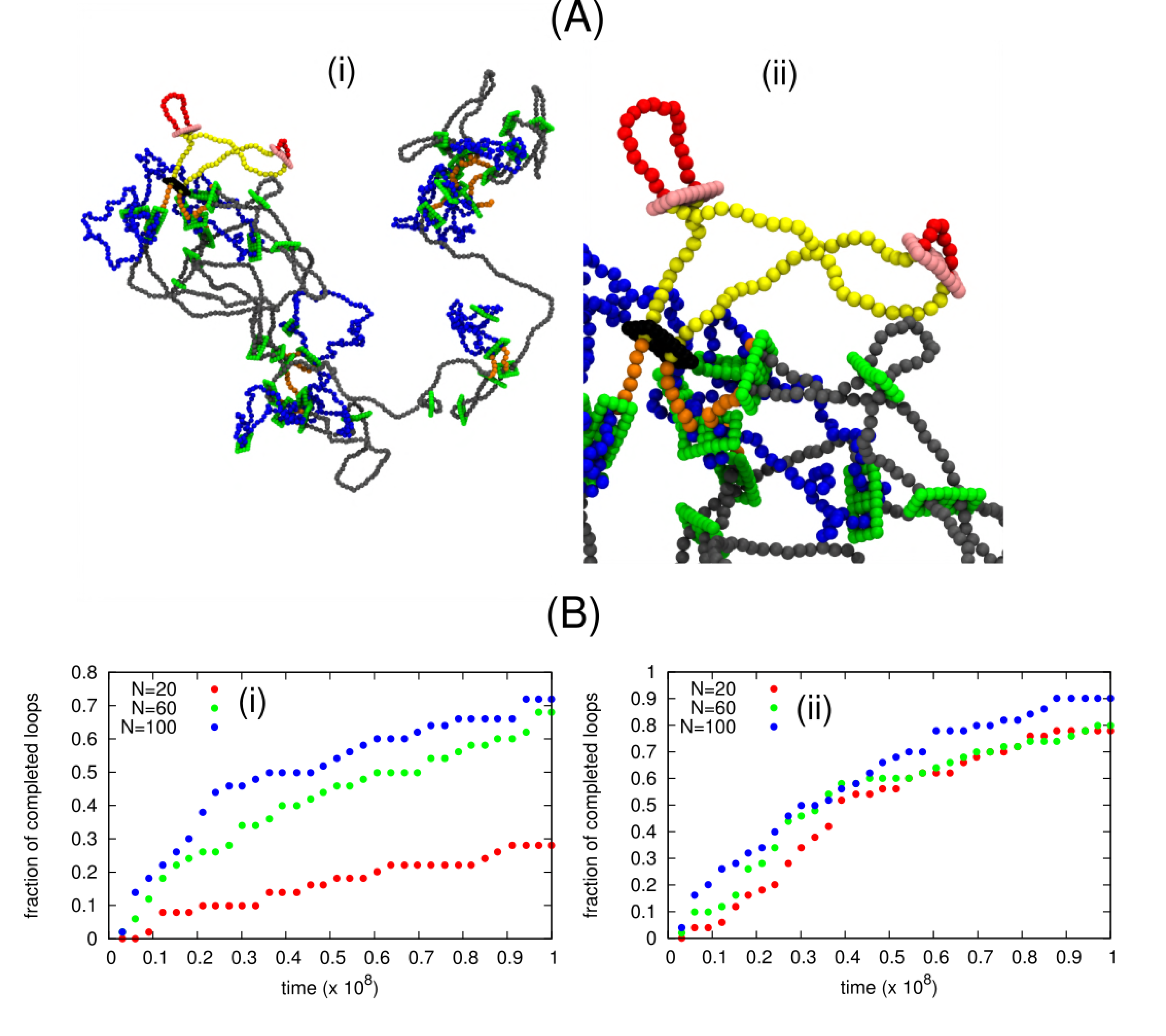
Results from 3-D simulations considering different numbers of slip-links. (A) Snapshots of the copolymer simulations, with flexible and stiff regions in the chromatin fibres shown in blue and grey respectively. (ii) shows a zoom of the snapshot in (i), illustrating the formation of nested and consecutive loops. The black slip-link forms the yellow loop, to which the consecutive red loops (formed by the pink slip links) are nested. Such configurations increase the efficiency of diffusive loop extrusion to form large loops. The ratcheting effect is in operation in the example shown in (ii); this is a stiff region, where nested loops are more visible, however it occurs in both stiff and flexible regions. (B) Percentage of completed CTCF loop (size as in Fig. 3) which are completed in simulations with *N* = 20, *N* = 60 or *N* = 100 slip-links (i.e., 1, 3 or 5 slip-link per section), for flexible [(i), *l*_*p*_ = 2*σ*] or stiff [(ii), *l*_*p*_ = 10*σ*] chromatin regions.

Our results for one slip-link per domain are consistent with those of Figure 2, suggesting that, as expected intuitively, the slip-link dynamics only depends on the *local* chromatin flexibility (rather than on global chromatin structure), at least these simulations which consider dilute chromatin. More interestingly, we find that the number of slip-links play an important role. While the general qualitative trend that diffusive sliding is faster on stiffer fibres remains true whatever the copy number, we find that the effect is quantitatively reduced as the number of slip-links increases (Figs. 5B). This is because the presence of multiple slip-links leads to a ratchet effect, first discussed in Ref. [20]. This effect arises as the slip-link-mediated loops are likely to be nested or stacked within each other (see highlighted snapshot in Fig. 5A). As a result, diffusive shrinkage of the largest (outer) loop is hindered by the presence of smaller (inner) loops, due to the steric repulsion between slip-links. Whilst the ratchet effect speeds up the growth of loops in all cases, it does so more strongly in the flexible polymer case, so that the gap between flexible and stiff loop formation efficiency narrows (Figs. 5B).

Interestingly, the ratchet effect is maximised when the loader is close to the CTCF binding sites. Indeed, additional simulations with the loader in the middle of each chromatin section (i.e., in between, and maximally distant from, CTCF binding sites), suggest that under the conditions considered here, increasing *N decreases* loop size. Thus, considering a copolymer as in Figure 5 with 1, 3 and 5 slip-links per CTCF section, and a loader in the middle of each, leads to a fraction of completed loops (after 10^6^ Brownian times) equal to respectively, 0.08 and 0.07 for *l*_*p*_ = 2*σ*, and to 0.83, 0.55 and 0.43 for *l*_*p*_ = 10*σ*. Our previous work in Ref. [20] found the ratchet to be efficient also when the loader was placed in the middle – the difference is presumably due to the fact that the simulations here consider a substantially smaller CTCF domain size. A possible reason for the increased efficiency of the ratchet effect when the loader is positioned close to the CTCF is that this setup may render less likely configurations where all loops are consecutive (as opposed to nested).

## Discussion and Conclusions

In summary, here we have used computer simulations to study the dynamics of diffusive loop extrusion by means of molecular slip-links on chromatin fibres of different flexibility.

Flexibility is potentially an important parameter which can vary along mammalian chro-mosomes. The common view is that active regions containing promoters, enhancers and transcribed regions are associated with open chromatin, which is more flexible with respect to that of inactive regions [31]. Recent microscopy work *in vivo* has also shown that the local thickness of the chromatin fibre and its density varies throughout the nucleus [32], and these changes are likely to be associated to a change in flexibility.

We have found that the diffusive motion of slip-links, similar to cohesins, is strongly affected by the flexibility of the underlying chromatin fibre. In particular, we have quantified the effect on diffusive loop extrusion – i.e., the creation of large chromatin loops via diffusive sliding. Whilst such chromatin loops would grow or shrink in the absence of other interactions, assuming that CTCF binds to cohesin in a directionality-dependent manner is sufficient to stabilise these loops, thereby rendering diffusive loop extrusion an appealing model to explain the formation of convergent CTCF loops in mammalian genomes. We have found here that diffusive loop extrusion is substantially faster and more efficient in stiff chromatin, which may be associated with heterochromatin rich in H1, but not in HP1 or other bridges (which would render it more compact and globular). Our results therefore suggest that cohesin may be an important player to compactify inert chromatin regions, where other chromatin bridges are depleted.

We have also shown that, when cohesin loading (mediated *in vivo* by proteins such as NIPBL) occurs preferentially near CTCF binding sites (which are also associated with DNase hypersensitive sites [33]), the simultaneous presence of multiple cohesin within the same stretch of chromatin further enhances the efficiency of diffusive loop extrusion.

Besides being relevant to our understanding of the fundamental mechanisms underlying chromatin looping and 3-D chromosome organisation, our results could potentially be tested in single-molecule setups with reconstituted chromatin fibres. We also hope they will be of use in designing more sophisticated simulations of chromatin folding, addressing for instance the interplay between molecular slip links such as cohesins and other transcription factors, which can also organise chromatin (see, e.g., [34, 35]).

## Author contributions

C.A.B., J.J., D. Michieletto, and D. Marenduzzo designed the research, performed the research, analyzed the data, and wrote the article.

## Acknowledgements

This work was supported by ERC (CoG 648050, THREEDCELL-PHYSICS).

## References

[1] J. Dekker, K. Rippe, M. Dekker and N. Kleckner. Capturing chromosome conformation. Science 295, 1306–1312 (2002).

[2] S. S. P. Rao, M. H. Huntley, N. C. Durand, E. K. Stamenova, I. D. Bochkov, J. T. Robinson, A. L. Sanborn, I. Machol, A. D. Omer, E. S. Lander et al. A 3D map of the human genome at kilobase resolution reveals principles of chromatin looping. Cell 159, 1665–1680 (2014).

[3] J. R. Dixon, S. Selvaraj, F. Yue, A. Kim, Y. Li, Y. Shen, H. Hu, J. S. Liu and B. Ren. Topological domains in mammalian genomes identified by analysis of chromatin interactions. Nature 485, (2012) 376-380.

[4] J. E, Phillips, E. Jennifer and V. G. Corces. CTCF: Master Weaver of the Genome. Cell 7, 1194–1211 (2009).

[5] Z. Tang et al. CTCF-mediated human 3D genome architecture reveals chromatin topology for transcription, Cell 163, 1611 (2015).

[6] M. Oti, J. Falck, M. A. Huynen and H. Zhou. CTCF-mediated chromatin loops enclose inducible gene regulatory domains. BMC Genomics 17, 252 (2016).

[7] K. Nasmyth. Cohesin: a catenase with separate entry and exit gates? Nat. Cell Biol. 13, 1170–1177 (2011).

[8] M. T. Ocampo-Hafalla and F. Uhlmann. Cohesin loading and sliding. J. Cell. Sci. 124, 685– 691 (2011).

[9] P. J. Huis in ‘t Veld, F. Herzog, R. Ladurner, I. F. Davidson, S. Piric, E. Kreidl, V. Bhaskara, R. Aebersold, and J. M. Peters. Characterization of a DNA exit gate in the human cohesin ring. Science 346, 968–972 (2014).

[10] F. Uhlmann. SMC complexes: from DNA to chromosomes. Nat. Rev. Mol. Cell. Biol. 17, 399–412 (2016).

[11] M. H. Kagey et al. Mediator and cohesin connect gene expression and chromatin architecture. Nature 467, 430 (2010).

[12] G. Fudenberg, M. Imakaev, C. Lu, A. Goloborodko, N. Abdennur, and L. D. Mirny. Formation of Chromosomal Domains by Loop Extrusion. Cell Reports 15, 2038–2049 (2016).

[13] A. L. Sanborn et al. Chromatin extrusion explains key features of loop and domain formation in wild-type and engineered genomes. Proc. Natl. Acad. Sci. USA 112, E6456 (2015).

[14] E. Alipour and J. F. Marko. Self-organization of domain structures by DNA-loop-extruding enzymes. Nucleic Acids Res. 40, 11202–11212 (2012).

[15] K. Nasmyth. Disseminating the genome: joining, resolving, and separating sister chromatids during mitosis and meiosis. Ann. Rev. Genet. 35, 673 (2001).

[16] T. Terakawa, S. Bisht, J. M. Eeftens, C. Dekker, C. H. Haering and E. C. Greene. The condensin complex is a mechanochemical motor that translocates along DNA. Science 358, 672–676 (2017).

[17] J. Stigler, G. Çamdere, D. E. Koshland and E. C. Greene. Single-Molecule Imaging Reveals a Collapsed Conformational State for DNA-Bound Cohesin. Cell Rep. 15, 988 (2016).

[18] I. F. Davidson et al. Rapid movement and transcriptional re-localization of human cohesin on DNA. EMBO J. 35, 2671 (2016).

[19] M. Kanke, E. Tahara, P. J. Huis in’t Veld, and T. Nishiyama. Cohesin acetylation and Wapl-Pds5 oppositely regulate translocation of cohesin along DNA. EMBO J. 35, 2686–2698 (2016).

[20] C. A. Brackley, J. Johnson, D. Michieletto, A. N. Morozov, M. Nicodemi, P. R. Cook and D. Marenduzzo. Nonequilibrium chromosome looping via molecular slip links. Phys. Rev. Lett. 119, 138101 (2017).

[21] C. A. Brackley, J. Johnson, D. Michieletto, A. N. Morozov, M. Nicodemi, P. R. Cook and D. Marenduzzo. Extrusion without a motor: a new take on the loop extrusion model of genome organization. Nucleus, https://doi.org/10.1080/19491034.2017.1421825.

[22] D. Racko, F. Benedetti, J. Dorier and A. Stasiak. Transcription-induced supercoiling as the driving force of chromatin loop extrusion during formation of TADs in interphase chromosomes. Nucl. Acids Res., gkx1123, https://doi.org/10.1093/nar/gkx1123.

[23] C. R. Calladine, H. R. Drew, B. F. Luisi and A. A. Travers, Understanding DNA, 3rd Edition, Elsevier Academic Press, London. (2004).

[24] B. Alberts et al., Molecular biology of the cell, Garland Science, New York (2002).

[25] J. Langowski. Polymer models of DNA and chromatin. Eur. Phys. J. E 19, 241 (2006).

[26] A. N. Boettiger, B. Bintu, J. R. Moffitt, S. Wang, B. J. Beliveau, G. Fudenberg, M. Imakaev, L. A. Mirny, C. Wu, and X. Zhuang, Super-resolution imaging reveals distinct chromatin folding for different epigenetic states, Nature, 529, 7586 (2016).

[27] T. Sexton, E. Yaffe, E. Kenigsberg, F. Bantignies, B. Leblanc, M. Hoichman, H. Parrinello, A. Tanay, and G. Cavalli, Three-dimensional folding and functional organization principles of the Drosophila genome, Cell, 148, 3 (2012).

[28] H. Hajjoul, J. Mathon, H. Ranchon, I. Goiffon, J. Mozziconacci, B. Albert, P. Carrivain, J. M. Victor, O. Gadal, K. Bystricky and A. Bancaud. High-throughput chromatin motion tracking in living yeast reveals the flexibility of the fiber throughout the genome. Genome Res. 23, 1829 (2013).

[29] A. Rosa and R. Everaers. Structure and Dynamics of Interphase Chromosomes. PLOS Comp. Biol., e1000153 (2008).

[30] L. Ringrose, S. Chabanis, P. Angrand, C. Woodroofe and A. F. Stewart. Quantitative comparison of DNA looping in vitro and in vivo: chromatin increases effective DNA flexibility at short distances, EMBO J. 18, 6630–6641 (1999).

[31] P. R. Cook and D. Marenduzzo. Entropic organization of interphase chromosomes. J. Cell Biol. 186, 825–834 (2009).

[32] H. D. Ou, S. Phan, T. J. Deerinck, A. Thor, M. H. Ellisman and C. O’Shea. ChromEMT: Visualizing 3D chromatin structure and compaction in interphase and mitotic cells. Science 357, eaag0025 (2017).

[33] Y.-M. Wang, P. Zhou, L.-Y. Wang, Z.-H. Li, Y.-N. Zhang and Y.-X. Zhang. Correlation Between DNase I Hypersensitive Site Distribution and Gene Expression in HeLa S3 Cells. PLoS ONE 7, e42414 (2012).

[34] M. Barbieri, M. Chotalia, J. Fraser, L.-M. Lavitas, J. Dostie, A. Pombo and M. Nicodemi. Complexity of chromatin folding is captured by the strings and binders switch model. Proc. Natl. Acad. Sci. USA 109, 16173–16178 (2012).

[35] C. A. Brackley, J. Johnson, S. Kelly, P. R. Cook and D. Marenduzzo. Simulated binding of transcription factors to active and inactive regions folds human chromosomes into loops, rosettes and topological domains. Nucleic Acics Res. 8, 3503 (2016).

